# A chromosome-level genome assembly of a vernal pool specialist amphibian, the Western Spadefoot, *Spea hammondii*

**DOI:** 10.1101/2025.11.16.688715

**Authors:** Ben Thompsky, Eric Beraut, Robert D. Cooper, Merly Escalona, Robert E. Espinoza, Robert N. Fisher, Courtney Miller, Oanh Nguyen, Samuel Sacco, Ruta Sahasrabudhe, William E. Seligmann, Erin Tofflemier, Ian J. Wang, H. Bradley Shaffer

## Abstract

We assembled and annotated a chromosome-level reference genome for the Western Spadefoot, *Spea hammondii* (Anura, Scaphiopodidae) representing one of only three amphibians included in the California Conservation Genomics Project (CCGP). *Spea hammondii* is a vernal pool breeding anuran native to California and northwestern Baja California which has undergone both range contractions and local extirpations across its distribution, primarily due to habitat loss and degradation and drought. The species is recognized by the state of California as a Species of Special Concern and is proposed for listing under the United States Endangered Species Act.

Using the established CCGP pipeline, this *S. hammondii* genome was produced using Pacific Biosciences HiFi long-reads and Omni-C proximity ligation, resulting in a *de novo* genome assembly 1.14 Gb in length, distributed across 479 scaffolds (scaffold N50 = 120.8 Mb; largest scaffold = 183.6 Mb) with a BUSCO completeness score of 90.9% using a conserved tetrapod ortholog set. Our assembly shows high base accuracy (QV = 63.7) and low frameshift error in coding regions (QV 50.42). Annotation of this genome yielded 20,434 genes with a BUSCO completeness score of 94.7%. This reference genome, in combination with range-wide resequencing data from CCGP, will facilitate statewide population genomic assessments to delineate conservation units, quantify inbreeding and genomic load, and test for adaptive variation associated with vernal pool hydrology and drought tolerance, all of which are important considerations in the proposed federal listing.

## Introduction

The Western Spadefoot, *Spea hammondii*, is a species of medium-sized fossorial anuran that was historically widespread in the low-elevation, non-desert regions of central and southern California and northern Baja California, Mexico (Fig. 1). The species often occurs in discrete and isolated populations tied to small seasonal wetlands, including human-constructed livestock ponds. Adult *S. hammondii* are primarily found in grassland, chaparral and oak woodland habitats with loose soil conducive to burrowing (Morey 2005; Hansen and Shedd 2025). During the non-rainy season, post-metamorphic juveniles and adults live underground in self-constructed burrows, preferring sand and silt over clay soils (Baumberger 2019). They emerge from burrows during rainy nights, generally in the winter and spring. *Spea hammondii* typically breed between January and June. Breeding is wholly dependent on the presence of ephemeral water bodies, rendering the species exceptionally vulnerable to drought and climatic changes that lower or change the frequency of precipitation (Thomson *et al*. 2016).

**Figure 1.**
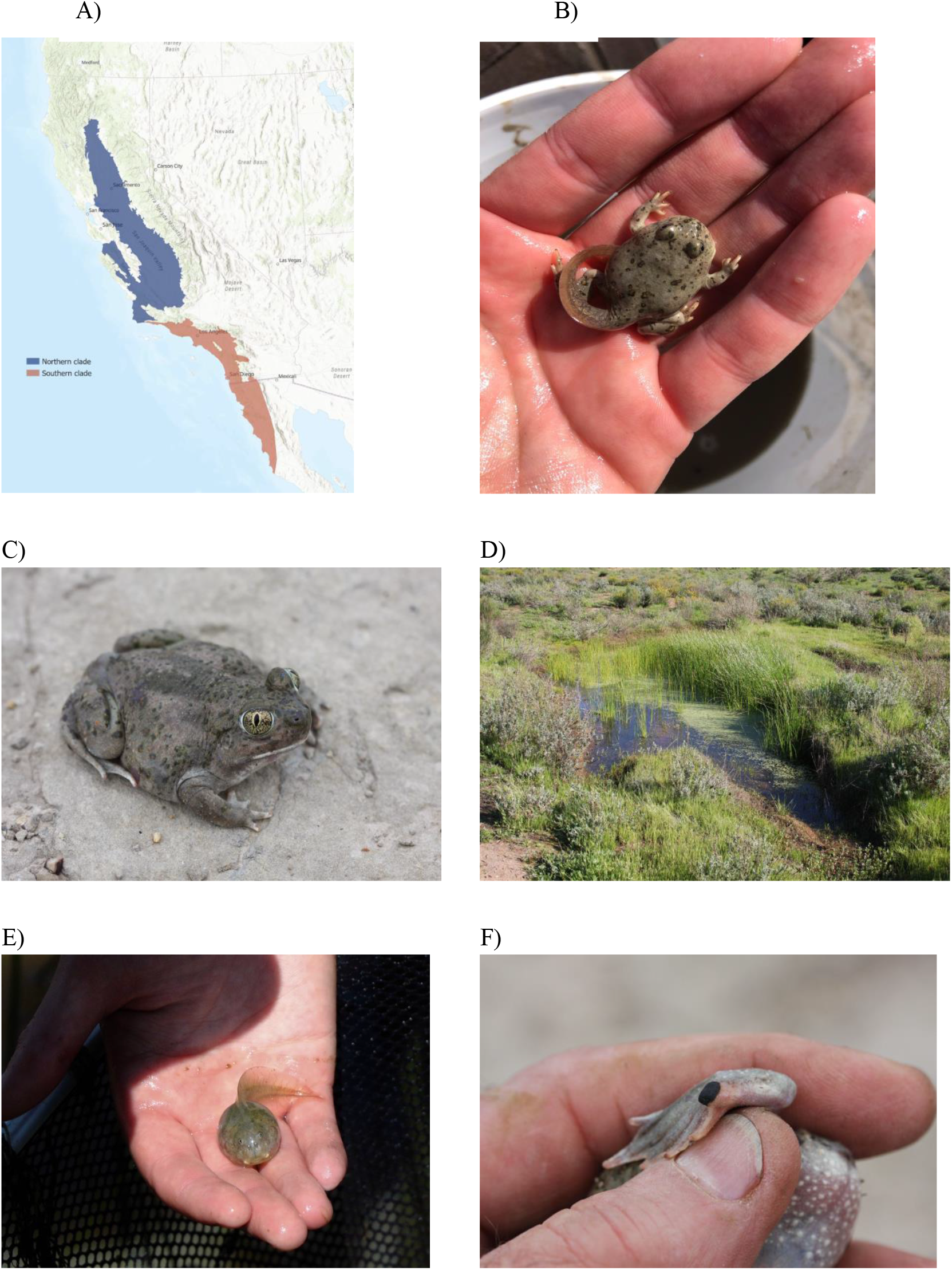
A) Range map of *Spea hammondii* showing the two genetically separate subpopulations north and south of the Transverse Ranges as well as range extension into Baja California(Kristi Cripe [CDFW], map). B) Mid-metamorphosis spadefoot tadpole. Animals at this stage have well-developed spades on their rear feet and are capable of completing metamorphosis on land (Robert D. Cooper, image). C) An adult Western spadefoot from northern Santa Barbara County, California (H. Bradley Shaffer, image). D) Vernal pool used by *S. hammondii* for breeding. This pool is located in San Diego County, California (H. Bradley Shaffer, image). (E) *Spea hammondii* tadpole, displaying their relatively large size (Nora Papian/USFWS, Public Domain, https://www.fws.gov/media/spadefoot-toad-tadpole). F) Close-up of the keratinized spade on the rear foot of an adult *S. hammondii*. Spadefoots use this spade to dig burrows in loose sandy soil (H. Bradley Shaffer, image)

Larval development is extremely rapid and has been the focus of substantial research on *S. hammondii*. In short-hydroperiod vernal pools, *S. hammondii* exhibit rapid development, with smaller sizes at metamorphosis. Larvae also show remarkable life history adaptations and phenotypic plasticity, with the potential to develop into cannibalistic carnivorous tadpoles with increased keratinization of the jaw in response to prey density and pond dry-down (Levis *et al*. 2018). While this ‘cannibal morph’ phenotype is better documented in the other three species in the genus (Levis *et al*. 2015), at least one published observation from northwestern Baja California has documented cannibalistic behaviors in metamorphosing *S. hammondii* (Alvarez *et al*. 2024).

The taxonomic history of the genus *Spea*, including S. *hammondi*, has been a source of confusion at multiple levels. Long placed in the family Pelobatidae, current taxonomy recognizes the New-World spadefoots as members of Scaphiopodidae, a small clade of seven species and two genera (*Spea* and *Scaphiopus*) (see Frost 2025 for a detailed taxonomic history). Scaphiopodids, along with the Old-World families Pelodytidae, Megophryidae and Pelobatidae, form a clade that is the sister group to Neobatrachia, a large clade that includes well over 90% of living anurans (Pyron and Wiens 2011). Recent genetic analyses (Neal *et al*. 2018) has identified two species-level lineages that are divided into northern and southern groups by the Transverse Ranges, a mid-elevation mountain range that often forms a biogeographic boundary between many northern and southern California biotic elements (Rissler *et al*. 2006). Neal *et al*. (2018) suggested that each clade should be managed as distinct conservation units, based on both genetic data and species distribution modeling. While the single range-wide entity is currently recognized by California as a Species of Special Concern (Thomson *et al*. 2016) the northern and southern clades are each currently under consideration for listing under the United States Endangered Species Act (USFWS 2023).

This genome release is part of the California Conservation Genomics Project (CCGP), which provides genomic resources and their analyses to inform conservation actions at a statewide scale (Shaffer *et al*. 2022; Fiedler *et al*. 2022). This chromosome-level reference genome, the third available for the genus (see also *S. multiplicata* (Seidl *et al*. 2019), GenBank accession GCA_009364415.1, *S. bombifrons*, GenBank accession GCF_027358695.1) provides a key resource to help address outstanding questions in the systematics and conservation of *S. hammondii*. As part of the CCGP, this genome will enable range-wide population-genomic analyses to delineate conservation units, quantify inbreeding and genomic load, and map adaptive variation associated with hydroperiod, temperature, and drought (Shaffer *et al*. 2022). This information is of particular importance for this species as it exists in increasingly disjointed and potentially inbred populations, making a reference genome vital to develop cost-effective marker panels for genetically informed reintroductions, translocations, and long-term monitoring across California’s fragmented seasonal-wetland landscapes. In congeneric *Spea*, reference genomes have provided a framework for understanding and locating the metabolic and endocrine pathways implicated in phenotypic plasticity (Seidl *et al*. 2019); this genome may provide the ability to determine whether these adaptations are also present in *S. hammondii*. Finally, this reference genome adds to a growing set of California vernal pool specialist species, including the freshwater crustaceans *Branchinecta lynchi* (Blair *et al*. 2023a), *Branchinecta lindahli* (Blair *et al*. 2023b), and *Lepidurus packardi* (Blair *et al*. 2022), and the vernal pool plants *Neostapfia colusana* (Pennington *et al*. 2025) and *Tuctoria greenei* (Toews *et al*. 2025), that will allow cross-taxon analyses of the genomic health of California’s rich, endemic, and extremely endangered vernal pool ecosystems.

## Methods

### Biological materials

We sequenced an adult *S. hammondii* (HBS 135690) from the southern clade/distinct population segment (Neal *et al. 2018)* that was collected as a tadpole from a drying ephemeral puddle (a tire track) on a dirt trail in Los Angeles county (Coordinates: 34.4680837, 118.5479482) and reared in captivity for approximately 2.5 yrs in the California State University Northridge vivarium and euthanized on 19 August 2020. Liver, thigh muscle, tongue, and heart tissues were harvested and flash-frozen in liquid nitrogen within two min of euthanasia, and the remaining carcass was formalin-fixed for later museum deposition.

### High molecular weight genomic DNA isolation

High molecular weight genomic DNA (HMW gDNA) was isolated from a liver sample. Flash frozen liver tissue (20 mg) was homogenized using a mortar and a pestle in liquid nitrogen. Homogenized tissue was lysed with 2ml of lysis buffer containing 100 mM NaCl, 10 mM Tris-HCL-pH 8.0, 25 mM EDTA, 0.5% SDS and 100 µg/ml proteinase K overnight at room temperature. Lysate was treated with 20 µg/ml RNase at 37 C for 30 min and cleaned with equal volumes of phenol/chloroform using phase-lock gels (Quantabio, Beverly, MA, USA; Cat # 2302830). DNA was precipitated by adding 0.4X volume of 5M ammonium acetate and 3X volume of ice-cold ethanol. The resulting DNA pellet was washed twice with 70% ethanol and resuspended in an elution buffer (10mM Tris, pH 8.0). Purity of DNA was assessed with a spectrophotometer (NanoDrop ND-1000, Thermo Fisher Scientific, Waltham, MA, USA) where 260/280 ratios of 1.88 and 260/230 of 2.14 were observed. Total DNA yield of 13.3 µg was obtained as quantified using Qbit 2.0 Fluorometer (Thermo Fisher Scientific, Waltham, MA, USA). DNA fragment size was quantified using a Femto pulse system (Agilent Technologies, Santa Clara, CA, USA); 44% of the DNA was in fragments > 100 kilobases (kb).

### HiFi library preparation

The HiFi SMRTbell library was constructed using the SMRTbell Express Template Prep Kit v2.0 (Pacific Biosciences - PacBio, Menlo Park, CA, USA; Cat. #100-938-900) according to the manufacturer’s instructions. HMW gDNA was sheared to a target DNA size distribution between 15 – 20 kb using Diagenode’s Megaruptor 3 system (Diagenode, Belgium; Cat. B06010003). The sheared gDNA was concentrated using 0.45X of AMPure PB beads (PacBio, Cat. #100-265-900) for the removal of single-strand overhangs at 37°C for 15 min followed by further enzymatic steps of DNA damage repair at 37°C for 30 min, end repair and A-tailing at 20° C for 10 minand 65° C for 30 min, ligation of overhang adapters v3 at 20° C for 60 min and 65° C for 10 min to inactivate the ligase, then nuclease treated at 37° C for 1 hour. The SMRTbell library was purified and concentrated with 0.45X AMPure PB beads for size selection using the BluePippin/PippinHT system (Sage Science, Beverly, MA, USA; Cat #BLF7510/HPE7510) to collect fragments greater than 7-9 kb. The 15 – 20 kb average HiFi SMRTbell library was sequenced at the UC Davis DNA Technologies Core (Davis, CA, USA) using two 8M SMRT cell (PacBio, Cat #101-389-001), Sequel II sequencing chemistry 2.0, and 30-h movies each on a PacBio Sequel IIe sequencer.

### Omni-C library preparation

The Omni-C library was prepared using the Dovetail Omni-C Kit (Dovetail Genomics, Scotts Valley, CA, USA) according to the manufacturer’s protocol with slight modifications. First, liver tissue was thoroughly ground with a mortar and pestle while cooled with liquid nitrogen. Subsequently, chromatin was fixed in place in the nucleus. The suspended chromatin solution was then passed through 100 μm and 40 μm cell strainers to remove large debris. Fixed chromatin was digested under various conditions of DNase I until a suitable fragment length distribution of DNA molecules was obtained. Chromatin ends were repaired and ligated to a biotinylated bridge adapter followed by proximity ligation of adapter-containing ends. After proximity ligation, crosslinks were reversed and the DNA was purified from proteins. Purified DNA was treated to remove biotin that was not internal to ligated fragments. An NGS library was generated using an NEB Ultra II DNA Library Prep kit (New England Biolabs, Ipswich, MA, USA) with an Illumina compatible y-adaptor. Biotin-containing fragments were then captured using streptavidin beads. The post-capture product was split into two replicates prior to PCR enrichment to preserve library complexity with each replicate receiving unique dual indices. The library was sequenced at Vincent J. Coates Genomics Sequencing Lab (Berkeley, CA, USA) on an Illumina NovaSeq 6000 platform (Illumina, San Diego, CA, USA) to generate approximately 100 million 2 x 150 bp read pairs per giga base pair (Gb) of genome size.

### Transcriptome RNA extraction, library preparation, and sequencing

Total RNA from 3 tissues (tongue, muscle, heart) was extracted using a Qiagen RNeasy Mini Kit (Qiagen, Netherlands), according to the manufacturer’s protocol. RNA libraries were then prepared using the KAPA mRNA HyperPrep Kit (Roche, Switzerland) according to the manufacturer’s protocol. Libraries were sequenced with 100bp reads on an Illumina NovaSeq 6000 platform (Illumina), to generate approximately 50M reads per library at the Technology Center for Genomics and Bioformatics (UCLA, CA, USA) on an Illumina NovaSeq X.

### Nuclear genome assembly

We assembled the genome of a *S. hammondii* individual following the CCGP assembly pipeline Version 5.0, as outlined in Table 1, where we list the tools and non-default parameters used in the assembly process. We removed the remnants adapter sequences from the PacBio HiFi dataset using HiFiAdapterFilt (Sim *et al*. 2022) and generated an initial diploid phased assembly using HiFiasm (Cheng *et al*. 2021) in HiC mode with the filtered PacBio HiFi reads and the Omni-C short-reads, a process that generates two assemblies, one per haplotype. We then aligned the Omni-C data to each assembly following the Arima Genomics Mapping Pipeline (https://github.com/ArimaGenomics/mapping_pipeline) and scaffolded them independently with SALSA (Ghurye *et al*. 2017, 2019).

**Table 1.**
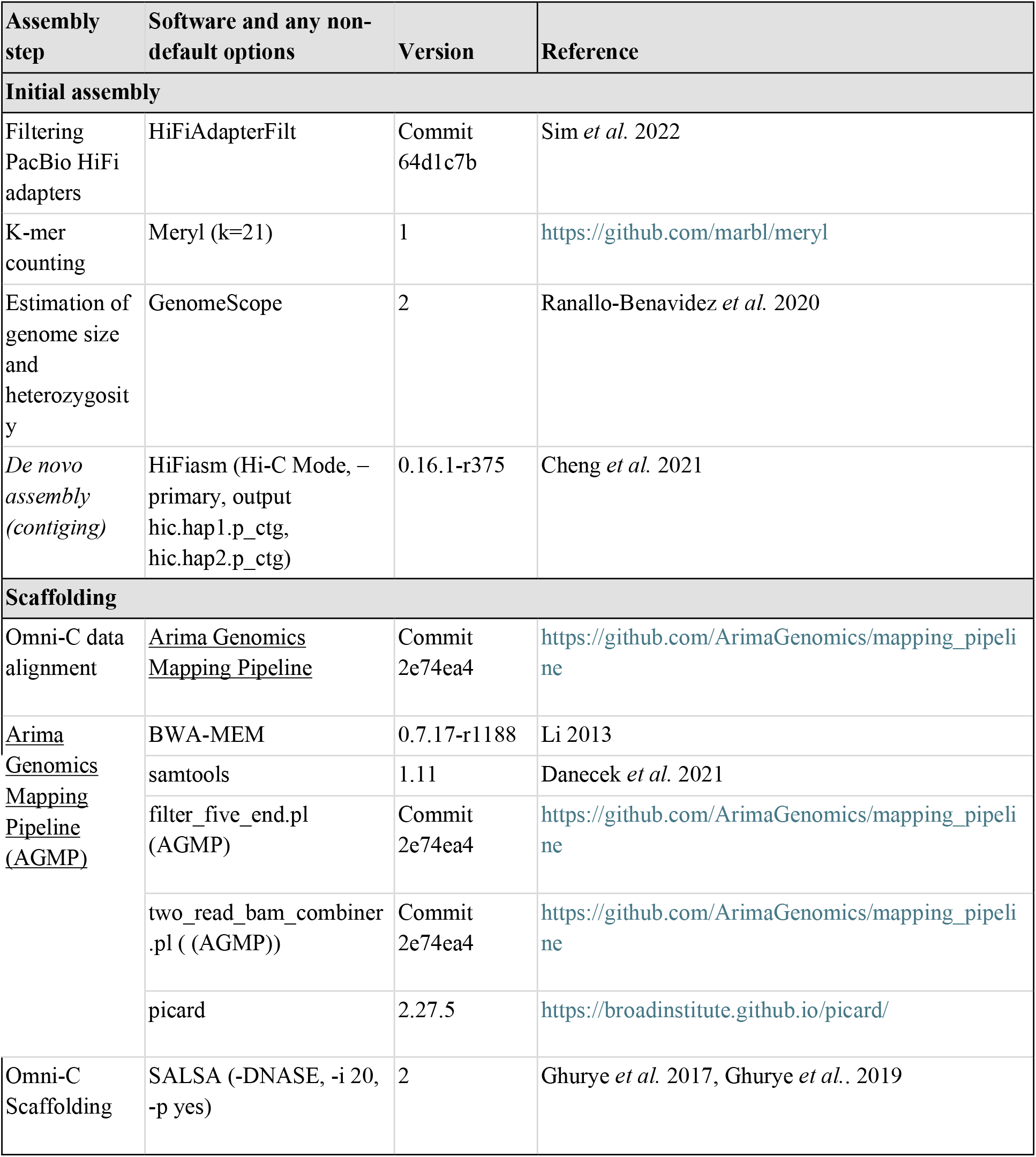
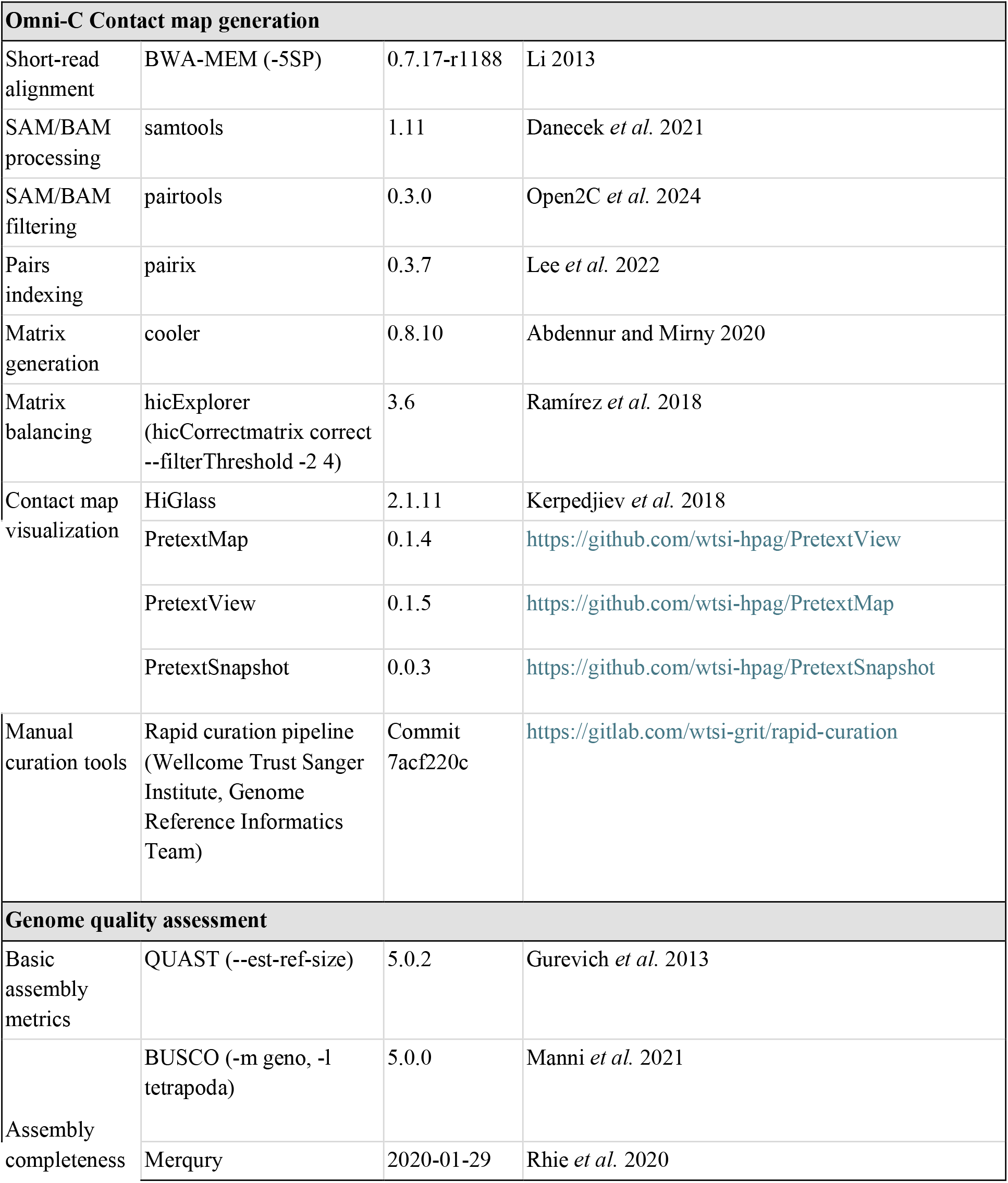
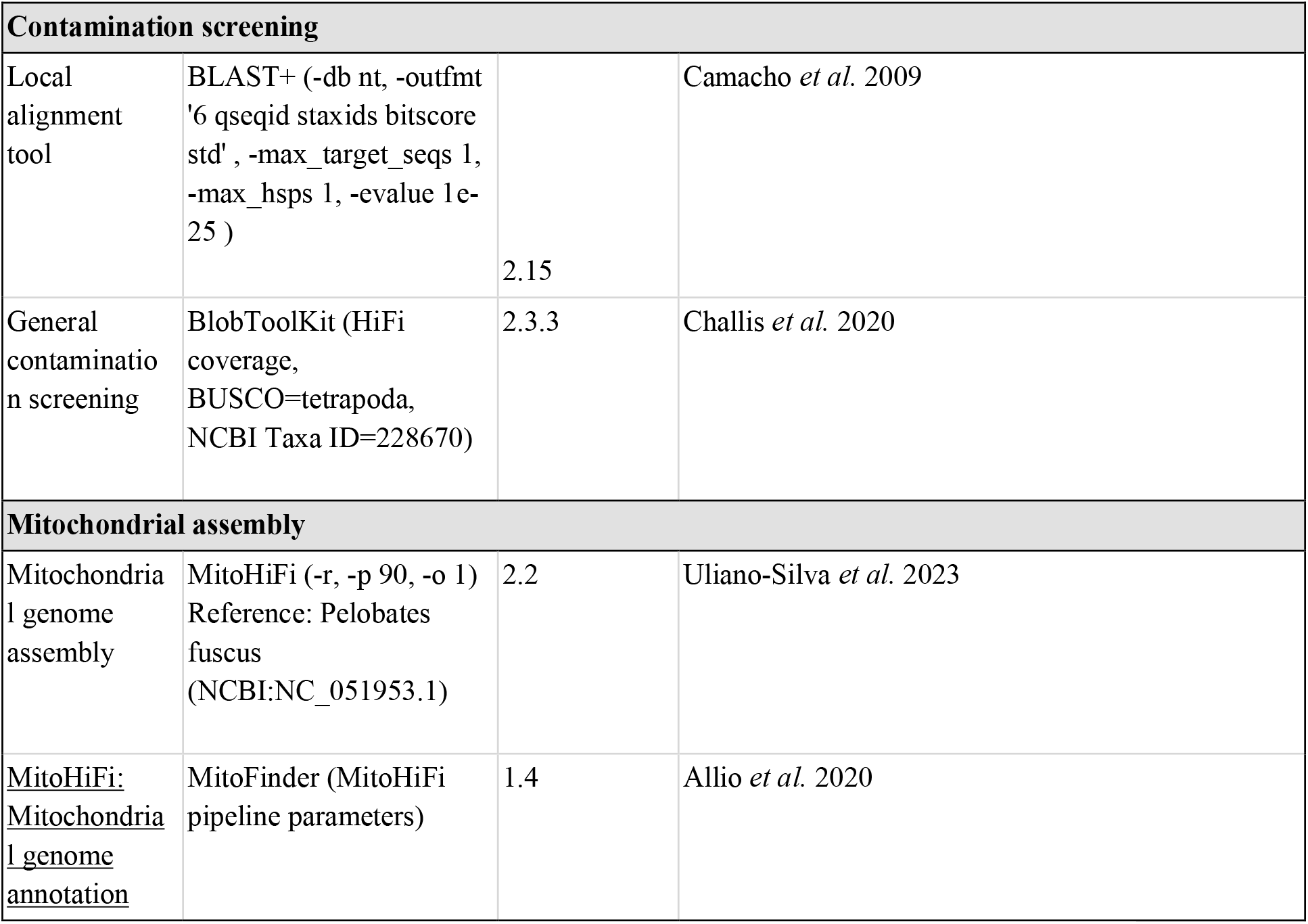
Assembly pipeline and software used.

The assemblies for both haplotypes were manually curated by iteratively generating and analyzing their corresponding Omni-C contact maps. To generate the contact maps, we aligned the Omni-C data with BWA-MEM (Li 2013), identified ligation junctions, and generated Omni-C pairs (Lee *et al*. 2022) using pairtools (Open2C *et al*. 2023). Then, we generated multi-resolution Omni-C matrices with cooler (Abdennur and Mirny 2020) and balanced them with hicExplorer (Ramírez *et al*. 2018). We used HiGlass (Kerpedjiev *et al*. 2018) and the PretextSuite (https://github.com/wtsi-hpag/PretextView; https://github.com/wtsi-hpag/PretextMap; https://github.com/wtsi-hpag/PretextSnapshot) to visualize the contact maps. We identified misassemblies and misjoins in these contact maps and modified the assemblies using the Rapid Curation pipeline from the Wellcome Trust Sanger Institute, Genome Reference Informatics Team (https://gitlab.com/wtsi-grit/rapid-curation). Some of the remaining gaps (joins generated during scaffolding and/or curation) were closed using the PacBio HiFi reads and YAGCloser (https://github.com/merlyescalona/yagcloser). We checked for contamination using the BlobToolKit Framework (Challis *et al*. 2020).

### Mitochondrial genome

We assembled the mitochondrial genome of *S. hammondii* from the PacBio HiFi reads using the reference-guided pipeline MitoHiFi (Allio *et al*. 2020; Uliano-Silva *et al*. 2023). The mitochondrial sequence of a *Pelobates fuscus* (NCBI:NC_051953.1) was used as the starting sequence. After completion of the nuclear genome, we searched for matches of the resulting mitochondrial assembly sequence in the nuclear genome assembly using BLAST+ (Camacho *et al*. 2009) and filtered out contigs and scaffolds from the nuclear genome with a percentage of sequence identity > 99% and size smaller than the mitochondrial assembly sequence.

### Genome quality assessment and size estimation

We generated k-mer counts from the PacBio HiFi reads using meryl (https://github.com/marbl/meryl). The k-mer counts were then used in GenomeScope2.0 (Ranallo-Benavidez *et* 2020) to estimate genome features including genome size, heterozygosity, and repeat content. To obtain general contiguity metrics, we ran QUAST (Gurevich et*t* 2013). To evaluate genome quality and functional completeness, we used BUSCO (Manni *et al*. 2021) with the Tetrapoda ortholog database (tetrapoda_odb10) which contains 5,310 genes. Assessment of base level accuracy (QV) and k-mer completeness were performed using the previously generated meryl database and merqury (Rhie *et al*. 2020). We further estimated genome assembly accuracy via BUSCO gene set frameshift analysis using the pipeline described in Korlach *et al*. (2017). Measurements of the size of the phased blocks were based on the size of the contigs generated by HiFiasm on HiC mode. We follow the quality metric nomenclature established by Rhie *et al*. (2021), with the genome quality code x.y.P.Q.C, where, x = log10[contig NG50]; y = log10[scaffold NG50]; P = log10 [phased block NG50]; Q = Phred base accuracy QV (quality value); and C = % genome represented by the first ‘n’ scaffolds, following a karyotype of 2n=26 for this species (Wasserman 1970). Quality metrics for the notation were calculated on the assembly for the haplotype one assembly.

### Genome annotation

We annotated the genome assembly using the NCBI Eukaryotic Genome Annotation Pipeline v0.4.1-alpha (hereafter, “EGAPx”) which is published in the NCBI RefSeq database (O’Leary *et al*. 2016) and accessible through the NCBI GitHub page (“ncbi/egapx”). Annotation features were identified by aligning transcripts and proteins from related taxa in the RefSeq database using BLAST (Camacho *et al*. 2009). Novel, species-specific RNA-Seq reads were also aligned to the assembly using the alignment software STAR (Dobin *et al*. 2013). Additional features are predicted using HMM-based gene models using the NCBI Gnomon software. We evaluated the quality and completeness of our annotation by comparing the longest protein for each annotated coding gene to tetrapods (odb10) using BUSCO v5.7.1 (Manni *et al*. 2021). We report the number of annotation features and the BUSCO results in Table 2.

**Table 2.**
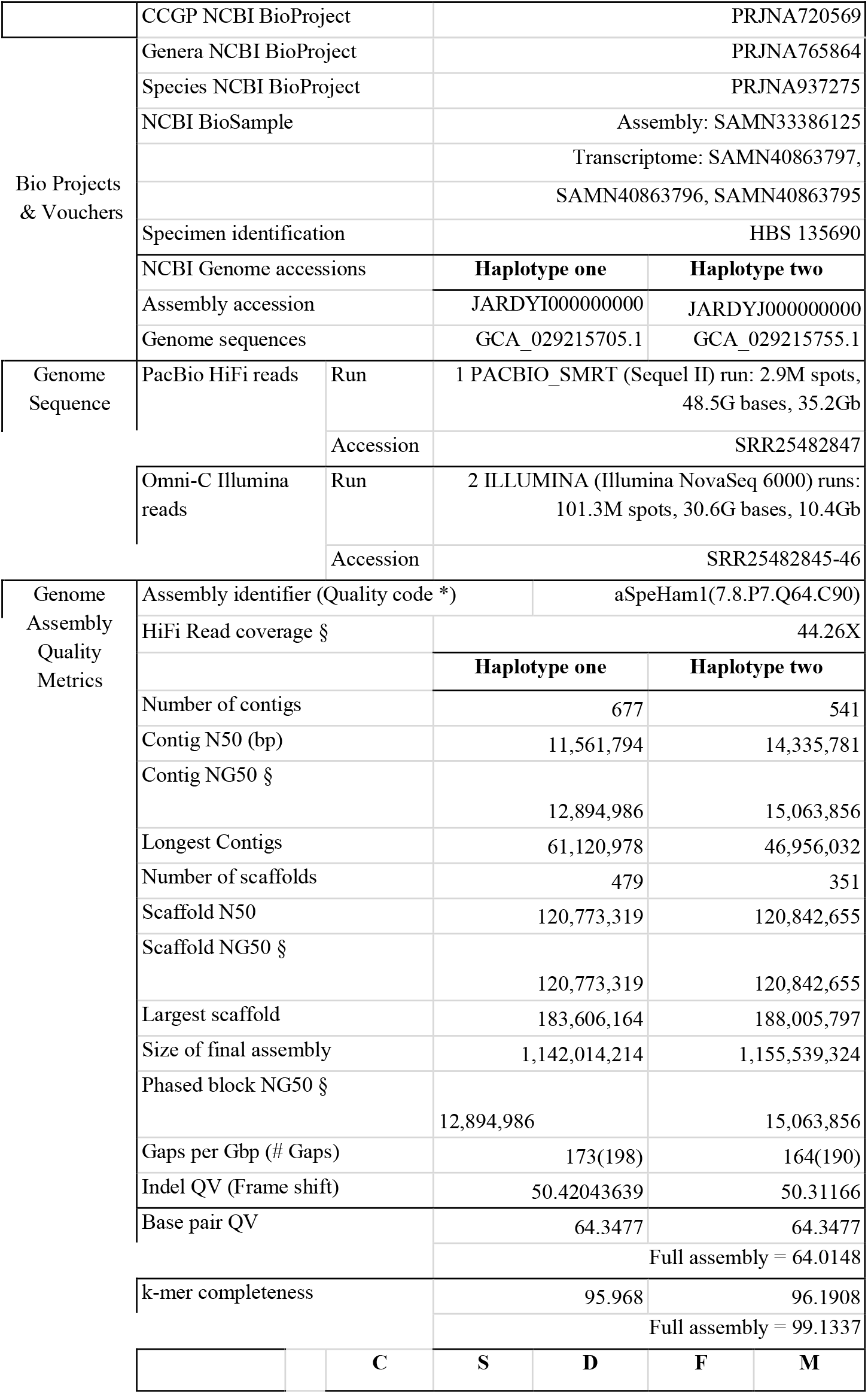
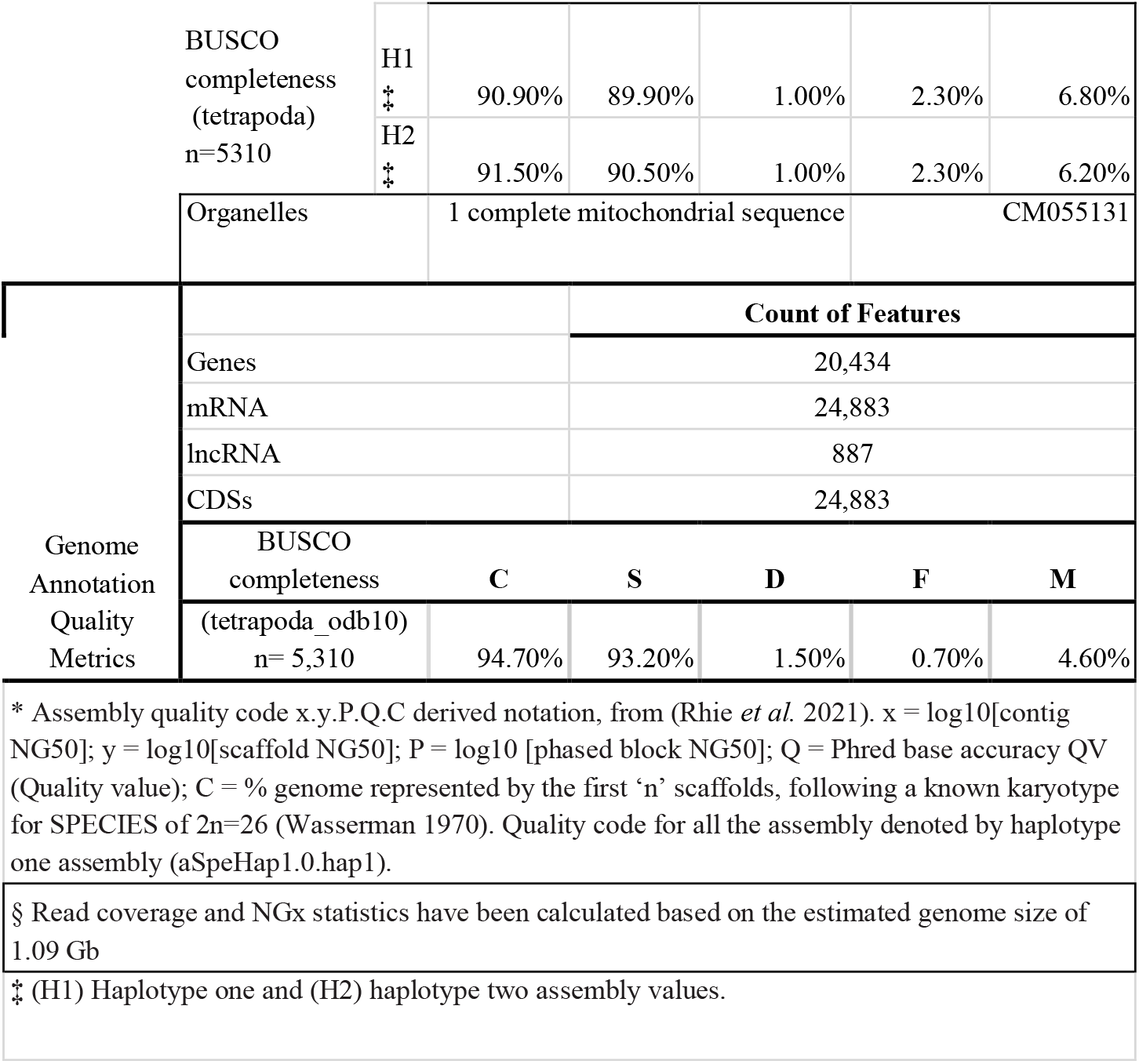
Sequencing and assembly statistics and accession numbers.

## Results

### Sequencing data

The Omni-C library generated 289.96 million read pairs, and the PacBio HiFi library generated 4.65 million reads. The PacBio HiFi sequences yielded ∼56X genome coverage and had an N50 read length of 18,307 bp; a minimum read length of 1 bp; a mean read length of 18,118 bp; and a maximum read length of 65,001 bp (see Supplementary Fig. S1 for read length distribution). Based on the PacBio HiFi data, GenomeScope 2.0 estimated a genome size of 1.09 Gb, a 0.164 % sequencing error rate, and 0.355% heterozygosity. The k-mer spectrum shows a unimodal distribution with a major coverage peak at ∼43-fold coverage (Figure 2A). Sequencing of mRNA libraries for SAMN40863797, SAMN40863796, and SAMN40863795 yielded 28M, 28M, 32M paired end reads, respectively.

**Figure 2.**
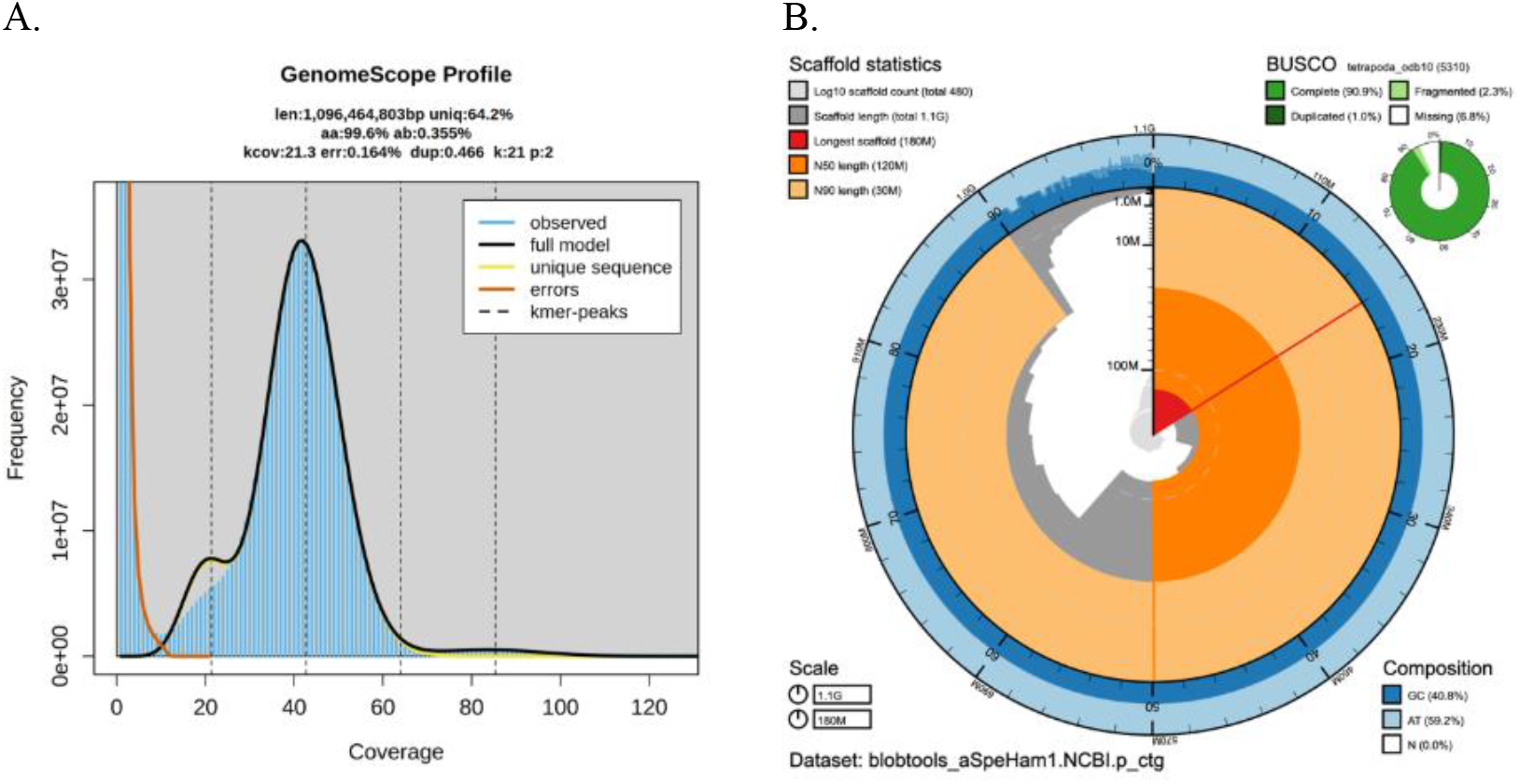
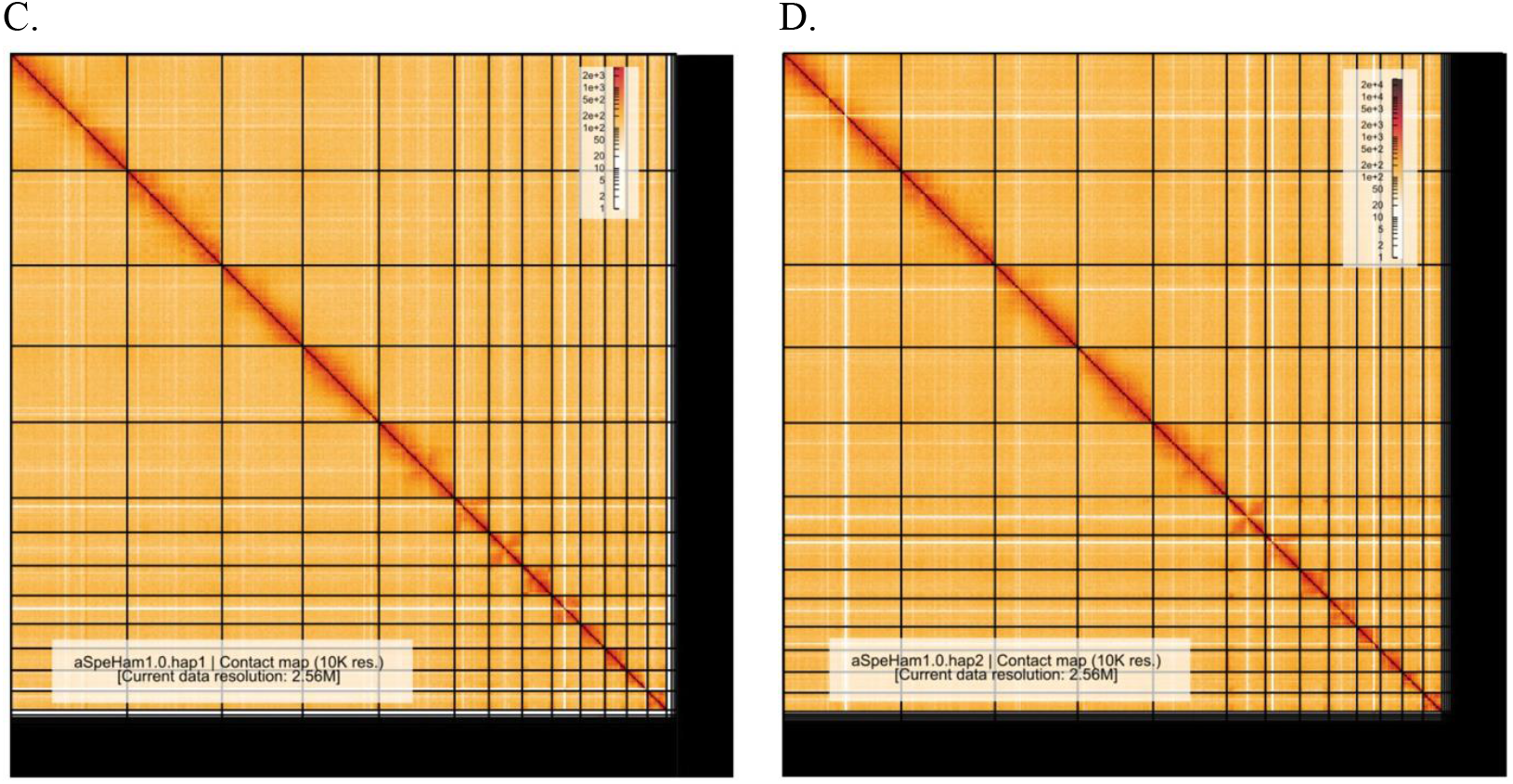
Visual overview of assembly metrics. A) K-mer spectrum output generated from PacBio HiFi data without adapters using GenomeScope2.0. The unimodal pattern observed corresponds to a haploid genome. K-mers covered at lower coverage and low frequency correspond to differences between haplotypes, whereas the higher coverage and high frequency k-mers correspond to the similarities between haplotypes. B) BlobToolkit Snail plot showing a graphical representation of the quality metrics presented in Table 2 for the *Spea hammondii* primary assembly (haplotype one; GenBank accession no. GCA_029215705.1). The plot circle represents the full size of the assembly. From the inside-out, the central plot covers length related metrics. The red line represents the size of the longest scaffold; all other scaffolds are arranged in size-order moving clockwise around the plot and drawn in grey starting from the outside of the central plot. Dark and light orange arcs indicate the scaffold N50 and scaffold N90 values. The central light grey spiral shows the cumulative scaffold count with a white line deliniating each order of magnitude. White regions in this area reflect the proportion of Ns in the assembly. The dark vs. light blue area around it shows mean, maximum and minimum GS vs. AT content at 0.1% intervals (Challis *et al*. 2020). For BUSCO summary data, the definitions are ‘complete and single copy’ (Comp.) or ‘complete and duplicated’ (Dupl.). C-D) Omni-C Contact maps for the haplotype 1 (2C) and haplotype 2 (2D) assemblies generated with PretextSnapshot. Omni-C contact maps translate proximity of genomic regions in 3-D space to contiguous linear organization. Each line in the contact map corresponds to sequencing data supporting the linkage (or join) between two of such regions. Scaffolds are separated by black lines, and higher density corresponds to high levels of fragmentation.

### Nuclear genome assembly

The final genome assembly (aSpeHam1) consists of two phased haplotypes. Both assemblies are similar in size, but not equal to the estimated genome size from GenomeScope2.0, as has been observed in other taxa (*e*.*g*. Pflug *et al*. 2020).

The haplotype one assembly (aSpeHam1.0.hap1) consists of 479 scaffolds spanning 1.14 Gb with a contig N50 of 11.56 Mb, a scaffold N50 of 120.77 Mb, the largest contig size of 61.12 Mb, and the largest scaffold size of 183.6 Mb. The haplotype two assembly (aSpeHam1.0.hap2) consists of 351 scaffolds spanning 1.15 Gb with a contig N50 of 14.33 Mb, a scaffold N50 of 120.84 Mb, the largest contig size of 46.95 Mb, and the largest scaffold size of 188 Mb. The haplotype one assembly has a BUSCO completeness score for the Tetrapoda gene set of 90.9%, a base pair quality value (QV) of 63.7, a k-mer completeness of 95.96%, and a frameshift indel QV of 50.42. The haplotype two assembly has a BUSCO completeness score for the Tetrapoda gene set of 91.5%, a base pair QV of 64.35, a k-mer completeness of 96.1%, and a frameshift indel QV of 50.31.

During manual curation we made 217 joins (103 on haplotype one and 114 on haplotype two) and 28 breaks (12 on haplotype one and 16 on haplotype two) based on the Omni-C contact map signal. We closed 20 gaps (9 on haplotype one and 11 on haplotype two). We filtered out two contigs corresponding to mitochondrial contamination. No further contigs were removed. The Omni-C contact maps show highly contiguous assemblies supporting the genome organization of 13 chromosomes per haplotype (Fig. 2B). Classic cytogenetic work reports a diploid number of 2n=26 for *S. hammondii*, historically treated as *Scaphiopus hammondii* (Wasserman 1970). By sorting our haploid assembly scaffolds by length, we found that the largest 13 contain 90.87% of the total genome length. We believe the remaining sequence represents repeat-dense, unresolved regions and is likely contained in the smaller scaffolds. Assembly statistics are reported in Table 2 and represented graphically in Fig. 2C. We have deposited the genome assembly on NCBI GenBank (See Table 2 and Data Availability for details).

### Mitochondrial genome assembly

We assembled a partial mitochondrial genome for *S. hammondii* with MitoHiFi. The final mitochondrial sequence has a size of 16,682 bp, with a base composition of A=29.85%, C=27.89%, G=, 15.2%, T=27.07%, and consists of 2 rRNAs, 22 unique transfer RNAs, and 13 protein-coding genes.

### Genome Annotation

Our final genome annotation for *S. hammondii* included 20,434 genes, with a tetrapoda BUSCO completeness of 94.70%. A list of annotation statistics and BUSCO score breakdowns are reported in Table 2.

## Discussion

There are presently 63 chromosome-level genomes for anurans in the NCBI database, representing 42 species. To this data set, we add a chromosome-level reference genome for the Western Spadefoot, *S. hammondii*. Assembly quality is reflected in core metrics (N50 = 120.8 Mb) with the largest scaffolds reaching 183.6-188.0 Mb, and completeness (BUSCO = 90.9% against a conserved tetrapod ortholog set). Omni-C contact maps support long-range contiguity, and our scaffold N50 exceeds the expected mean chromosome length of pelobatoid frogs (∼88 Mb, using the pelobatoid haploid karyotype of 13), justifying the chromosomal scale of this assembly.

Amphibian genomes are remarkably variable in size, with many anurans showing clade-specific genome expansions or contractions. Compared to other anurans, *Spea* genomes are extremely compact. Of the 495 anuran genomes reported in the Animal Genome Size Database (https://www.genomesize.com), scaphiopodids are almost always among the smallest 10 species; their reported c-values (average = 1.42, n = 17 observations) are much smaller than anurans generally (average = 4.47, n = 495). Zeng *et al*. (2014) analyzed the genome size distribution within pelobatoid frogs (a group including *Spea* and related families) and found a strong statistical association between larval development time and genome size. However, the genome size statistics reported by Zeng *et al*. (2014) are also derived from c-values in the Animal Genome Size Database rather than assembled genomes, and these two estimates can vary. Our assembly spans 1.142 Gb, which aligns with the other two available *Spea* genomes but is on the low end of previously published c-values for *S. hammondii* (c-values of 1.27, 1.34, 1.55, and 2.34; Gregory 2025). We interpret this discrepancy as reflecting the modest underrepresentation of repetitive sequence in our current assembly and/or variability in older Feulgen c-value estimates. Regardless, as additional annotated genomes of anurans are produced, future studies should explore whether the developmental time/genome size correlation persists across pelobatoids and anurans more generally.

This *S. hammondii* reference genome also offers a comparative framework for studying phenotypic plasticity in spadefoots. Studies in congeners (*e*.*g. S. multiplicata*) found that rapid development and resource polyphenism involve shifts in lipid and cholesterol metabolism, and peroxisome biology (Levis *et al*. 2020). Our reference genome enables future mapping of these candidate pathways and their regulatory neighborhoods in *S. hammondii*, and testing for selection on these regions along hydroperiod gradients.

This reference genome is directly useful for supporting the conservation goals of the CCGP. Range-wide genome resequencing built on this coordinate system will enable delineation of conservation units across the patchy distribution for this species (including the genetic division north and south of the Transverse Ranges identified by Neal *et al*. [2018] based on RADseq data), estimation of relatedness, runs of homozygosity, and quantification of genomic load on a population basis. Landscape genetics analyses can identify loci linked to hydroperiod, desiccation tolerance, developmental rate, and temperature sensitivity. If inbreeding is detected, this information can be used to plan reintroductions and assisted gene flow to potentially minimize the negative effects of inbreeding while preventing the proliferation of harmful alleles and the degradation of natural diversity.

## Supplementary Material

Supplementary material is available at *Journal of Heredity* online.

## Acknowledgements

PacBio Sequel II/IIe library prep and sequencing were carried out at the DNA Technologies and Expression Analysis Core at the UC Davis Genome Center, supported by NIH Shared Instrumentation Grant 1S10OD010786-01. Deep sequencing of Omni-C libraries used the Novaseq S4 sequencing platforms at the Vincent J. Coates Genomics Sequencing Laboratory at UC Berkeley, supported by NIH S10 OD018174 Instrumentation Grant. We thank the staff at the UC Davis DNA Technologies and Expression Analysis Core and the UC Santa Cruz Paleogenomics Laboratory for their diligence and dedication to generating high quality sequence data. We thank the vivarium staff in the Department of Biology at California State University, Northridge for animal care and financial support from the College of Science and Mathematics. RNA extraction, library preparation and sequencing was carried out at the UCLA Technology Center for Genomics & Bioinformatics. We thank Robert Hansen at USGS for additional review of the manuscript as well as Kristi Cripe at CDFW for generating the range map in Fig. 1. Any use of trade, firm, or product names is for descriptive purposes only and does not imply endorsement by the U.S. Government.

## Data Availability

Data generated for this study are available under NCBI BioProject PRJNA937275. Raw sequencing data for sample HBS135690 (NCBI BioSample SAMN33386125) are deposited in the NCBI Short Read Archive (SRA) under SRR25482847 for PacBio HiFi sequencing data, and SRR25482845-46 for the Omni-C Illumina sequencing data. GenBank accessions for both primary and alternate assemblies are GCA_029215705.1 and GCA_029215755.1; and for genome sequences JARDYI000000000 and JARDYJ000000000. The GenBank genome assembly for the mitochondrial genome is CM055131. Data generated for the annotation in this study are available under NCBI BioProject PRJNA1013362. Raw RNA sequencing data (NCBI BioSamples SAMN40863797, SAMN40863796 and SAMN40863795) are deposited in the NCBI SRA [SRR29346342, SRR29346343, SRR29346344], respectively). Assembly scripts and other data for the analyses presented can be found at the following GitHub repository: www.github.com/ccgproject/ccgp_assembly

